# Immune cell-specific Cre reporter mouse labels a subset of CNS neurons

**DOI:** 10.1101/833194

**Authors:** Aurélie Bouteau, Botond Z. Igyártó

**Affiliations:** Thomas Jefferson University, Department of Microbiology and Immunology, Philadelphia, PA 19107

**Keywords:** Langerhans cells, langerin, brain, neuron, central nervous system

## Abstract

HuLangerin-Cre-YFP^f/f^ mice were generated to specifically mark a subset of antigen presenting immune cells, called Langerhans cells (LCs). During histological characterization of these mice, we found that, in addition to LCs an uncharacterized cell population in the central nervous system (CNS) also expressed YFP. In this study, we found that the CNS YFP^+^ cells were negative for microglia and astrocyte markers, but they expressed mature neuronal marker NeuN and showed neuronal localization/morphology. Thus, these mice might be used to study the ontogeny, migration and the role of a subset of CNS neurons.

## 1. Introduction

Langerhans cells (LCs) are localized to the epidermis and they constitute a subset of antigen presenting dendritic cells (DCs) (Clausen and Stoitzner, 2015). LCs have dual origins, yolk sac and fetal liver, and they share common precursors with microglia (Ginhoux and Merad, 2010). LCs participate in promotion of self-tolerance, anti-fungal immunity, skin immunosurveillance, and protective humoral immune responses (Kashem et al., 2017).

Langerin is a C-type lectin receptor that was originally described as a specific marker of LCs (Valladeau et al., 2000). Later studies have found that langerin is also expressed by other subsets of DCs (Bursch et al., 2007; Poulin et al., 2007). Langerin is required for the formation of Birbeck granules, tennis racquet-shaped cytoplasmic organelles, which were shown to play role in HIV degradation (de Witte et al., 2007). Langerin can recognize mannose and other sugars, but no internal signaling pathway associated with it has been described to date (Clausen and Stoitzner, 2015; Romani et al., 2010).

Human langerin driven Cre mice were generated to selectively activate or delete genes in LCs (Kaplan et al., 2007). Interestingly the human langerin promoter in mice appears to recapitulate the human expression profile (Welty et al., 2013). The huLangerin-Cre-YFP^f/f^ mice were shown to permanently label the LCs in the skin, mucosal surfaces and CD103/CD11b double positive DCs in the gut lamina propria, but not other mouse DCs (Welty et al., 2013).

In this manuscript we show that huLangerin-Cre-YFP^f/f^ mice selectively label a subset of neurons in the CNS.

## 2. Materials and Methods

### 2.1. Mice

huLangerin-Cre-YFP^f/f^ mice have been previously described (Kaplan et al., 2007). Rosa26 ^fl^STOP^fl^ DTA mice on C57BL/6 background were obtained from Jackson Laboratory. All experiments were performed with newborn to 12-week-old female and male mice. Mice were housed in SPF conditions in microisolator cages and fed irradiated food and acidified water. The Baylor Institutional Care and Use Committee approved all mouse protocols.

### 2.2. Histology

Adult and newborn pups were euthanized and their brains (including cerebellum and olfactory bulb) and spinal cords removed. The brain hemispheres were separated using surgical blades, immersed in buffered 1% PFA and fixed overnight in 4 °C refrigerator. After intensive washing in PBS the organ samples were infused with 15 % sucrose solution overnight and then frozen in OCT. The blocks were stored in −80 °C until use. Eight μm thick sagittal sections were prepared using a cryostat and dried at room temperature for 30 minutes. After quick rehydration in PBS the sections were stained in humidified chambers at room temperature for one hour with the following antibodies: anti-GFP/YFP-AF488 (Rockland), anti-GFAP (GA5; ThermoFisher Scientific), and anti-NeuN (A60; Millipore-Sigma). The sections were then rinsed in PBS, mounted with Aqua-Poly/Mount solution and imaged with a Nikon Ti microscope.

### 2.3. Flow Cytometry

Adult mice were euthanized and their brains (including cerebellum and olfactory bulb) collected, disrupted in a glass tissue grinder using RPMI (ThermoFisher Scientific) supplemented with DNAse (0.1 mg/mL; Sigma) and Liberase TL (0.3 Wunsch units/mL; Sigma). The disrupted tissue was then incubated in a CO_2_ incubator at 37 °C for 1 hour, with intermittent mixing. The resulting cell suspension was filtered through 40 μm cell strainer, washed with PBS, layered on Percoll gradient (30% and 70%) and centrifuged. The cells at the interphase of 30/70% layers were collected, washed in PBS and stained with the following antibodies from BioLegend: CD11b (M1/70), CD45.2 (104), F4/80 (BM8) and CX3CR1 (SA011F11) on ice for 30 minutes. All the flow cytometric plots presented in this manuscript were pre-gated on live (using Live/Dead stain) and singlet events. Samples were analyzed on LSRFortessa flow cytometer (BD Biosciences). Data were analyzed with FlowJo software (TreeStar).

## 3. Results

### 3.1. HuLangerin-Cre-YFP^f/f^ mice harbor YFP^+^ cells in the CNS

HuLangerin-Cre-YFP^f/f^ mice were made to track LCs, professional antigen presenting immune cells localized to the epithelia and lymph nodes (Kaplan et al., 2007). The mice were generated using BAC transgene that contains Cre under the regulation of human langerin promoter (Kaplan et al., 2007). In these mice the human langerin promoter drives the expression of Cre (Kaplan et al., 2007; Welty et al., 2013). Histological analysis revealed YFP^+^ cells in CNS of the huLangerin-Cre-YFP^f/f^ mice, but not in Cre^−^ littermate controls (**Fig. 1**). The YFP^+^ cells could be found throughout the CNS with certain regions of the brain, such as the hippocampus/dentate gyrus, cortical layers V-VI and some cerebral nuclei presenting with higher densities (**Fig. 1**). The YFP^+^ cells could be readily identified in newborn pups’ brain as well (histology not shown). Thus, these data suggest that the human langerin promoter becomes active sometimes during the embryogenesis, which allows the Cre-mediated recombination to take place and the YFP to be expressed.

**Figure 1.**
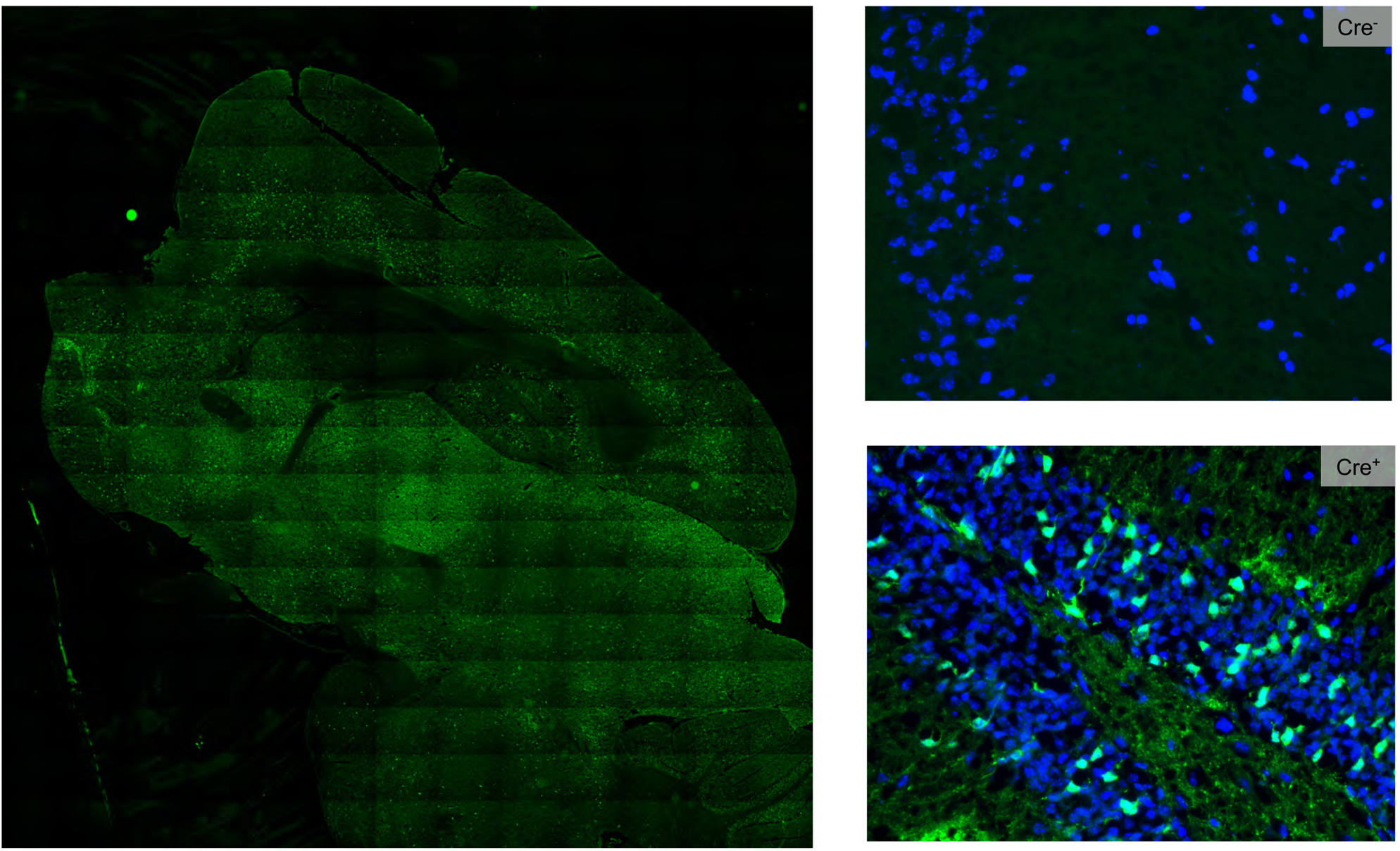
YFP^+^ cells in the CNS. Adult mouse CNS tissue from Cre^−^and Cre^+^ mice were fixed and stained with anti-YFP (green) and DAPI (blue). Left, stitched image of the Cre^+^ brain. Right lower, Cre^+^ dentate gyrus area of the CNS. Data from one representative experiment out of 10 is shown.

### 3.2. YFP^+^ cells are a subset of neurons

Since LCs share common precursors with microglia (Ginhoux et al., 2010; 2013; Hoeffel et al., 2012), we hypothesized that the YFP^+^ cells might be a subpopulation of microglia. To test this, we prepared single cell suspensions from adult brains and performed flow cytometry staining with classical mouse microglia markers: CD45, CD11b, F4/80 and CX3CR1 (Ginhoux et al., 2010; Jung, 2013; Waisman et al., 2015). We found that the YFP^+^ cells showed no reactivity with any of the microglia markers tested (**Fig. 2**). As next, we stained adult brain sections with astrocyte marker GFAP, and found no overlap with the YFP^+^ cells (**Fig. 3**). At last, all YFP^+^ cells showed staining with mature neuronal marker NeuN in different parts of CNS (**Fig. 4**). Thus, our data support that the YFP^+^ cells are a subset of neurons.

**Figure 2.**
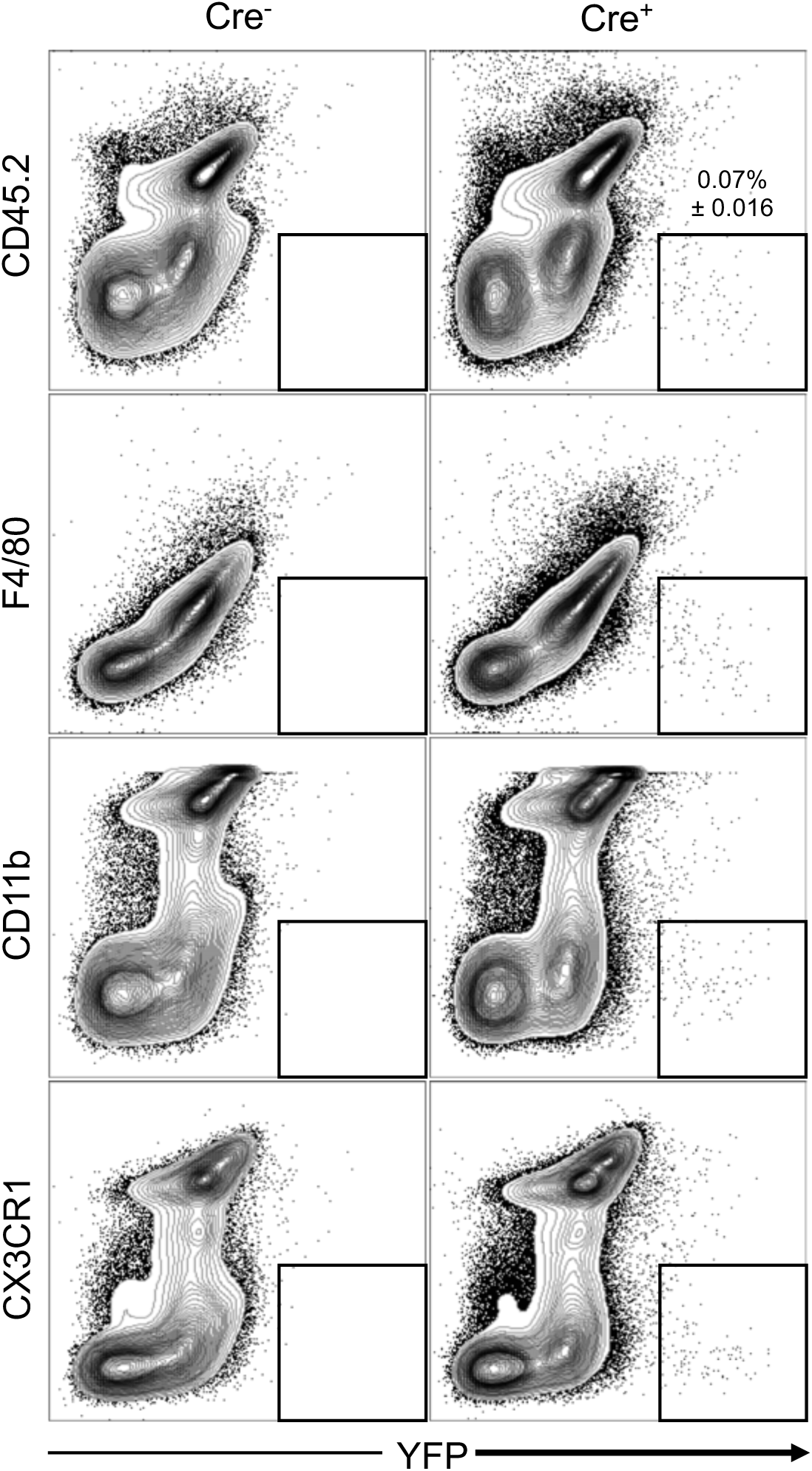
YFP^+^ cells do not express microglia markers. Cell suspensions prepared from the brains of Cre^−^and Cre^+^ mice were stained with the indicated markers and analyzed on flow cytometer. One representative experiment out of five is shown.

**Figure 3.**
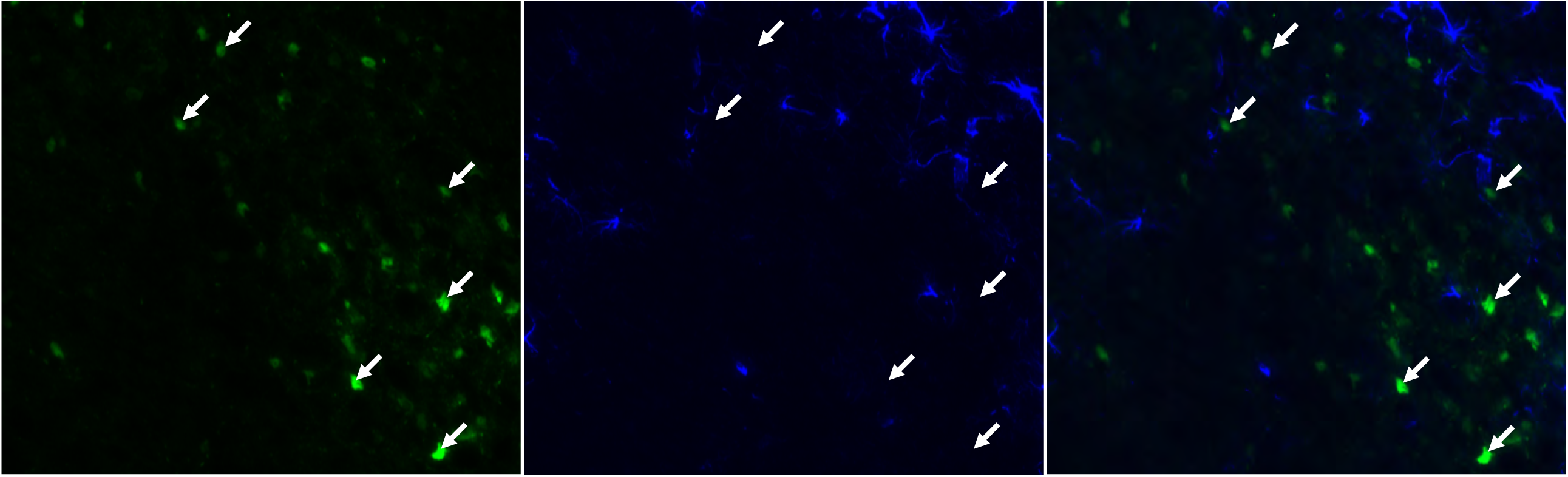
YFP^+^ cells do not express astrocyte marker GFAP. Adult mouse CNS tissue from Cre^+^ mice were stained with anti-YFP (green) and anti-GFAP (blue). Arrows indicate the localization of some of the YFP^+^ cells. Please note that there is no overlap between the YFP signal and GFAP.

**Figure 4.**
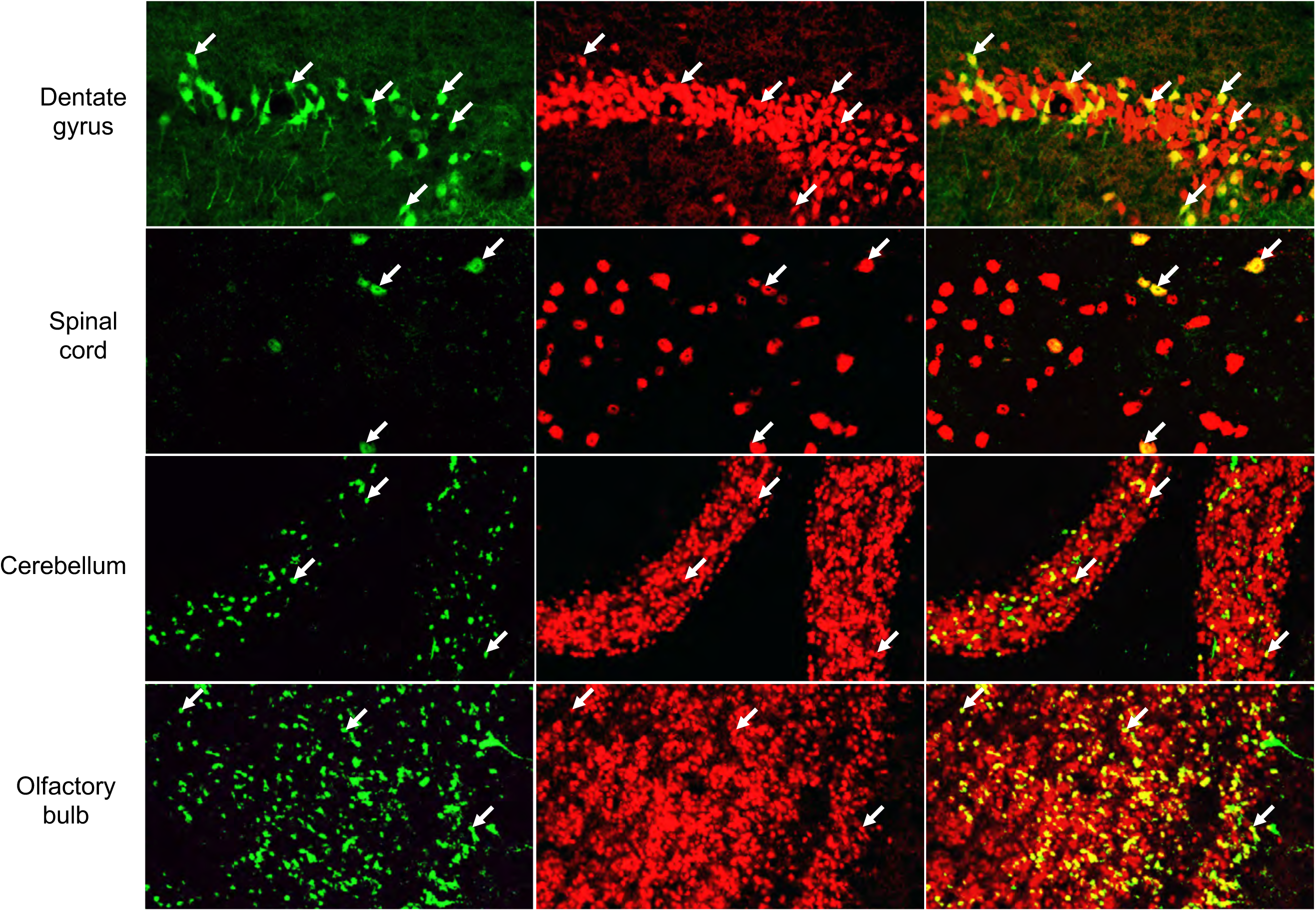
YFP^+^ cells express mature neuronal marker NeuN. Adult mouse CNS tissues from Cre^+^ mice were stained with anti-YFP (green) and NeuN (red). Arrows indicate the localization of some of the YFP^+^ cells. Please note the overlap between the YFP signal and NeuN.

## 4. Discussion

Here we show that huLangerin-Cre-YFP^f/f^ mice that were generated to track immune cells also label a subset of neurons in the CNS. The YFP^+^ cells were already present in the newborn’s CNS. The YFP^+^ neurons were present throughout the CNS with some regions, such as the olfactory bulb, cortical layers V and VI, hippocampal dentate gyrus, and cerebellum presenting with higher densities.

The presence of YFP^+^ cells in the newborns’ brain suggest that sometimes during the embryogenesis the human langerin promoter became active which led to Cre expression and permanent labeling of these cells with YFP. The human langerin transgenic mice appear to recapitulate the human pattern of langerin expression in the skin and the gut (Welty et al., 2013). Whether langerin is expressed in the human brain during embryonic development or after birth remains to be determined. Based on microarray data some low levels of langerin mRNA can be detected in different parts of the adult human brain (http://portal.brain-map.org/). Langerin is a C-type lectin receptor that recognizes different sugar moieties associated with microbes, however, no signaling pathways associated with langerin have been identified to date (Valladeau et al., 2003). The possible role if any of langerin expression in neurons remains to be determined.

The YFP^+^ cells are present in higher densities in the areas where adult neurogenesis takes place, such as the dentate gyrus (Gage and Temple, 2013; Homem et al., 2015). However, to determine whether new YFP^+^ neurons are also generated in adult brain a more careful time course analysis in combination with early neuronal markers, such as Sox2, DCX, and PSA-NCAM will be needed. The use of huLangerin-CreER^T2^-YFP^f/f^(Bobr et al., 2012) treated with tamoxifen at different stages of embryogenesis or adult life will also help to settle this question.

It will also be important to determine the nature of these neurons. Based on their widespread distribution in the CNS it is expected that the YFP^+^ cells to be a heterogenous population. Their heterogeneity might be due to different differentiation pathways that the YFP^+^ precursor cells have followed or because of distinct neuronal precursors were labeled with YFP. Their presence and localization in the dentate gyrus and cerebellum suggest that at least some of these YFP^+^ cells are granule cells. The YFP^+^ cells in the cortical layers V and VI could contribute to cortical efferent projections to basal ganglia, brain stem and spinal cord, and to the multiform or fusiform layer, projections to the thalamus, respectively. In the spinal cord they could be interneurons and/or part of the motoneuron system. Their identity could be confirmed by breeding these mice to different neuronal cell reporter mice and with the use of cell subset specific markers. A more detailed characterization could be achieved by RNA-seq performed on flow sorted cells or on samples isolated from different parts of the CNS using laser capture microscopy.

Our preliminary efforts to delete these cells by breeding the huLangerin-Cre-YFP^f/f^ mice to floxed “STOP” DTA mice, to gain insights into their physiological roles, remained unsuccessful. This could be due to the limitations of the Cre-lox system. It has been documented that when multiple floxed genes are combined with low Cre levels, preferential deletion of the certain targets could occur (Liu et al., 2013). These mice efficiently deleted the LCs in the skin and lymph nodes (data not shown). However, the YFP signal levels in the neurons are at least a log magnitude lower than in the LCs (unpublished observation). These data therefore suggest that in the neurons the Cre expression might not reach a level that would aid the deletion of the STOP codons on all the alleles and that could lead to non-toxic DTA levels.

In summary, here we describe a new mouse model that could contribute to our understanding of the development of the CNS, ontogeny, neurogenesis and neuronal migration patterns.

## Acknowledgments

We thank the animal facility, flow core and imaging core at BIIR for their help and support.

## Author contributions

B.Z.I. made the original observation. A.B. and B.Z.I. designed and performed the experiments. B.Z.I. wrote the manuscript. All authors participated in discussions of experimental results and edited the manuscript.

